# Zolpidem restores sleep and slows Alzheimer’s progression in a mouse model

**DOI:** 10.1101/2025.08.31.673377

**Authors:** Lu Yu, Shinya Yokomizo, Tri H. Doan, Qiuchen Zhao, Akshatha Ganne, Meenakshisundaram Balasubramaniam, Ksenia V. Kastanenka

## Abstract

**INTRODUCTION:** Deficits in Non-Rapid Eye Movement (NREM) sleep facilitate Alzheimer’s disease (AD) progression. Enhancing GABAergic signaling can restore sleep. Unbiased computational analysis identified zolpidem as high-affinity GABA receptor modulator facilitating chloride transport that could slow AD.

**METHODS:** Zolpidem’s effects on sleep and Alzheimer’s progression were evaluated in young APP/PS1 mice. Sleep was monitored with EEG/EMG telemetry. Widefield imaging with voltage-sensitive dyes was used to track sleep-dependent brain rhythms. Multiphoton microscopy allowed assessments of amyloid plaque load and basal neuronal calcium levels. Behavioral assays were used to measure memory and cognitive function.

**RESULTS:** Zolpidem restored NREM sleep and rescued sleep-dependent brain rhythm, slow oscillation. Zolpidem administration reduced cortical amyloid plaque burden, mitigated neuronal calcium overload, and enhanced sleep-dependent memory consolidation without adverse effects on locomotion.

**DISCUSSION:** Zolpidem effectively slowed Alzheimer’s progression in young APP/PS1 mice. This supports zolpidem’s therapeutic promise as an intervention strategy at early stages of AD.

## 1. BACKGROUND

Alzheimer’s disease (AD) is a progressive neurodegenerative disorder and the leading cause of dementia worldwide, affecting over 55 million people globally, a number expected to double by 2050 [1, 2]. It is clinically characterized by progressive cognitive decline and memory impairment. AD is pathologically characterized by the accumulation of amyloid-beta (Aβ) plaques and neurofibrillary tau tangles[3, 4].

Sleep impairments have emerged as a major modifiable risk factor for Alzheimer’s disease. Numerous studies have demonstrated that sleep deprivation and poor sleep quality are associated with increased amyloid-beta accumulation and accelerated disease progression[5, 6]. These pathological changes are closely linked to disruptions in the sleep-wake cycle, particularly deficits in non-rapid eye movement (NREM) sleep[7]. AD patients exhibit significant alterations in sleep architecture, including reduced NREM sleep duration, fragmented sleep, and diminished restorative sleep quality[8, 9]. Among these changes, impairments in slow oscillation (0.5-1 Hz), a sleep-dependent brain rhythm and the hallmark electrophysiological signature of deep NREM sleep, have gained increasing attention. Slow oscillation is a large-amplitude, low-frequency cortical rhythm that organizes synchronized neuronal activity across widespread brain networks. Each slow oscillation cycle consists of a down-state of relative neuronal silence and an up-state of synchronous neuronal firing[10–12]. Down and up states alternate during slow wave activity. Slow oscillation is important for restorative sleep, contributing to cognitive functions such as memory consolidation and learning in healthy individuals[10, 13]. Slow oscillation supports the homeostatic regulation of sleep, increasing in response to prolonged wakefulness and decreasing throughout the sleep period. In addition, slow oscillation helps organize faster brain rhythms, including sleep spindles and sharp-wave ripples, thereby enhancing the efficiency of neural networks involved in executive functions and synaptic plasticity[14–16]. Our published work showed that optogenetic rescue of slow waves slowed AD, while exacerbating slow wave deficits facilitated AD in a mouse model[17, 18]. Therefore, therapeutic targeting of slow oscillation could present a promising approach to slowing Alzheimer’s progression.

Mechanistically, impaired inhibitory neurotransmission has been identified as a major contributor to slow wave disruptions in young APP/PS1 (APP) mice, a mouse model of amyloidosis. Our published studies demonstrated that enhancing GABAergic tone through exogenous GABA administration restored slow oscillation in APP mice. However, pharmacological block of GABA receptors disrupted slow oscillation in healthy mice[18]. Similar to Alzheimer’s patients, young APP mice exhibit deficits in NREM sleep and increases in wake durations. Optogenetic stimulation of GABAergic interneurons in the anterior cortex effectively rescued sleep deficits and restored slow oscillation in APP mice [19]. These findings emphasize the importance of inhibitory circuits in maintaining healthy sleep rhythms and suggest that strategies aimed at enhancing inhibitory neurotransmission could have therapeutic benefits at early stages of AD.

Based on these insights, we performed an unbiased computational screen that identified zolpidem as high-affinity GABA_A_ receptor modulator facilitating chloride transport that could slow AD. We adopted a translational approach by testing zolpidem, an FDA-approved drug that enhances GABA_A_ receptor activity and promotes sleep. We systematically assessed the therapeutic potential of zolpidem in APP mice using multiple complementary experimental techniques. We first used in vivo voltage-sensitive dye imaging to directly visualize the acute effects of zolpidem on cortical slow oscillation dynamics across different doses, identifying 30 mg/kg as the optimal concentration for restoring slow oscillation. Wireless EEG/EMG telemetry was used to demonstrate that zolpidem improved NREM sleep architecture, enhancing NREM sleep continuity and stability while reducing sustained wakefulness. Importantly, chronic daily administration of zolpidem for four weeks significantly lowered amyloid plaque burden and neuronal calcium overload. Behavioral assessments revealed that zolpidem treatment selectively improved memory consolidation without adversely affecting locomotor activity or working memory. Collectively, our findings suggest that zolpidem, an FDA-approved agent, may serve as a promising therapeutic approach to restore sleep-dependent brain rhythms and ameliorate key pathological features of Alzheimer’s disease.

## 2. METHODS

### 2.1. Animals and Drug Administration

Male and female APPswe/PS1dE9 transgenic mice (APP, MMRRC 34829-JAX) were used. All mice were maintained under a 12:12 h light/dark cycle (7am lights on, 7pm lights off) with ad libitum access to food and water. For acute treatment experiments, APP mice aged 3–6 months were used to assess immediate effects of zolpidem on slow oscillation and sleep architecture, both of which were impaired during these ages[18–20]. For chronic treatment experiments, APP mice aged 7–9 months were used to evaluate the long-term impact of zolpidem on AD-related pathology, such as plaques and neuronal calcium overload, which are evident at these ages. Also, 7-9 month old APP mice were treated with zolpidem and followed with assessments of memory function, which is impaired at these ages[21]. All procedures were approved by the Institutional Animal Care and Use Committee (IACUC, protocol number 2012N000085) and conducted in accordance with NIH and ARRIVE guidelines.

### 2.2. Voltage-Sensitive Dye (VSD) Imaging

To evaluate the acute effects of zolpidem on cortical slow oscillation, wide-field in vivo imaging was performed using the voltage-sensitive dye RH2080. APP mice (3–6 months old) underwent craniotomy (5mm diameter) centered over the somatosensory cortex under 1–2% isoflurane anesthesia. RH2080 (1 mg/mL in PBS) was applied topically for 1 hour. After washing off excess dye with PBS and sealing the craniotomy with a glass coverslip, imaging was performed using a Hamamatsu ORCA-ER CCD camera (Hamamatsu Photonics) with a 630 ± 10 nm excitation filter and a >665 nm long-pass emission filter. Images were acquired at 20 Hz using Imaging Workbench software (INDEC Systems). A single dose of zolpidem (3,10, 30 mg/kg) or vehicle (0mg/kg) was administered via intraperitoneal (i.p.) injection, followed with repeated imaging at 5min intervals during the first 30 minutes post-injection, followed by longer intervals up to 120 minutes. 10% DMSO in PBS served as vehicle. Data were motion-corrected and spatially filtered before analysis. Slow oscillation power was extracted using temporal band-pass filtering (0.5–1 Hz) and quantified across the cortical field of view.

VSD image sequences were analyzed using ImageJ. For each recording, fluorescence changes were expressed as ΔF/F₀, where F₀ represented the minimum fluorescence intensity within the image sequence and ΔF was the change in pixel intensity relative to F₀. Mean ΔF/F₀ values were extracted from a manually defined region of interest (ROI) over the somatosensory cortex. To quantify oscillatory dynamics, Fourier transform analysis was performed in MATLAB, allowing calculation of oscillation power.

### 2.3. EEG/EMG Sleep Recordings

APP mice (3–6 months old) were surgically implanted with wireless EEG/EMG telemetry devices (HD-X02, Data Sciences International, Minneapolis, MN) under isoflurane anesthesia (3% induction, 1.5% maintenance) using aseptic techniques. Telemetry transmitters were placed subcutaneously along the dorsum. The skull was exposed and cleaned, and two stainless steel screws (M06-15-M-SS-P, US Micro Screw) serving as EEG electrodes were secured through the skull to contact the dura. The anterior electrode was placed 1 mm anterior to bregma and 1 mm lateral to the midline, while the posterior electrode was positioned 3 mm posterior to bregma and 3 mm contralateral to the anterior screw. EMG leads were sutured into the bilateral nuchal muscles. Following surgery, mice were allowed to recover for 10 days before recordings began. EEG/EMG signals were acquired using Ponemah Software v6.50 (DSI) and continuously recorded while animals were housed in individual home cages over receiver plates (RPC-1, DSI). Signals were digitized with a bandwidth of 0.1–200 Hz and sampled at 500 Hz. EEG/EMG data, activity, and temperature were recorded for 24 hours at baseline and for 48 hours following zolpidem (30 mg/kg, i.p.) or vehicle injection.

Sleep scoring was performed using NeuroScore Software v3.6 (DSI), which allowed fully automated sleep scoring to minimize experimenter bias as follows. Raw EEG and EMG signals were imported and bandpass filtered (EEG: 0.5–100 Hz; EMG: 10–100 Hz). Data were segmented into 10-second epochs and classified as wake, NREM sleep, or REM sleep using the integrated Mouse Sleep Scoring module. Wakefulness was defined by high-frequency, variable EEG activity and high EMG tone. NREM sleep was characterized by low-frequency, high-amplitude EEG and low EMG tone. REM sleep was identified by dominance of theta-band activity (theta/delta power ratio >3) and minimal EMG tone. Sleep bouts were defined as a minimum of two consecutive epochs of the same state. Power spectral density analysis was performed using fast Fourier transform (FFT) with a Hanning window applied to artifact-free epochs. Spectral bands were defined as follows: slow oscillation (0.5–1 Hz), delta (1–4 Hz), theta (4–8 Hz), alpha (8–12 Hz), sigma (12–16 Hz), and beta (16–24 Hz). Relative band power was computed as the ratio of individual band power to total power. Changes in sleep architecture and spectral composition were assessed across baseline and post-treatment periods.

### 2.4. Multiphoton Imaging of Amyloid and Calcium

To monitor AD-related pathology in vivo, we performed longitudinal two-photon imaging of amyloid plaques and neuritic calcium using Methoxy-X04 and the genetically encoded ratiometric calcium indicator Yellow Cameleon 3.6 (YC3.6) [22].

#### 2.4.1. AAV-YC3.6 Viral Injection

Mice were initially anesthetized with 5% isoflurane in oxygen and maintained at 1.5% isoflurane throughout the surgical procedure. Body temperature was maintained at approximately 37.5 °C using a heating pad, and ophthalmic ointment was applied to protect the eyes. Using sterile techniques, a 5 mm diameter round craniotomy was performed over the somatosensory cortex, alternating between the right and left hemispheres across animals to minimize lateralization bias.

To visualize basal neuronal calcium levels, AAV2-CBA-YC3.6 (2 µL, 2 × 10¹² vg/mL; University of Pennsylvania Vector Core) was injected bilaterally into the posterior somatosensory cortices of 7-month-old APP mice. Stereotaxic coordinates relative to bregma were: AP −1.5 mm, ML ±1.5 mm, DV −0.8 mm. The virus was delivered at a rate of 0.15 µL/min via a pulled glass micropipette, which was left in place for an additional 10 minutes post-injection to facilitate viral diffusion. Following viral injection, a sterile glass coverslip was positioned over the craniotomy and secured using dental cement to form a chronic cranial window. The skin was sutured around the window margin. Mice were allowed to recover on a heating pad until fully awake and mobile and were subsequently monitored daily. Experimental imaging began three weeks post-surgery to ensure complete recovery and sufficient transgene expression.

#### 2.4.2. Multiphoton Imaging of Amyloid Plaques

To visualize amyloid plaque pathology in vivo, two-photon imaging was performed through chronic cranial windows in mice. One day prior to imaging, mice received an intraperitoneal injection of Methoxy-X04 (10 mg/kg, Tocris), which binds to fibrillar amyloid and enables fluorescence-based detection. Immediately before imaging, Texas Red dextran (70 kDa, 2.5% in PBS; Thermo Fisher) was administered intravenously via the retro-orbital sinus to label the cerebral vasculature and facilitate anatomical orientation. Imaging was conducted using a FluoView FV1000MPE two-photon laser-scanning imaging system (Olympus) mounted on a BX61WI upright microscope with a 25× long working distance water-immersion objective (NA = 1.05, Olympus). A mode-locked Ti:Sapphire laser (MaiTai, Spectra-Physics) was tuned to 800 nm to excite Methoxy-X04. Images were acquired at 1× digital magnification, and Z-stacks were collected at 5 µm intervals. Body temperature was maintained using a heated stage, and mice remained under 0.5–1% isoflurane anesthesia during acquisition.

#### 2.4.3. Multiphoton Imaging of Calcium

To acquire basal neuronal calcium levels in APP mice, imaging of ratiometric YC3.6-expressing neuronal processes (neurites) was performed through the same cranial windows as imaging of amyloid plaques. APP mice were injected with AAV2-CBA-YC3.6 into the somatosensory cortex three weeks prior to imaging as described above (2.4.1). Imaging was performed on the same two-photon system using a Ti: Sapphire laser tuned to 860 nm to excite YC3.6. Fluorescence signals were separated using a 510 nm dichroic mirror and collected via PMTs set to detect CFP (380–480 nm) and YFP (500–540 nm). ROIs were selected based on morphology and expression clarity. Digital magnification was set to 2×. To minimize photodamage, laser power was maintained below 30 mW. Z-stack images were acquired at 1 µm intervals. Ratiometric (YFP/CFP) images were generated pixel-wise, and calcium-overloaded neurites were identified as those with mean YFP/CFP ratios exceeding 1.79, indicating cytosolic calcium levels >235 nM.

#### 2.4.4. Image Analysis

Z-stacks were aligned to baseline using rigid-body registration (TurboReg, ImageJ). Methoxy-X04–labeled amyloid plaques were segmented using intensity-based thresholding and reconstructed in 3D using Imaris (Bitplane). Total plaque burden was quantified as the sum of all segmented plaque volumes per field of view, and change over time was expressed as percent of baseline.

YC3.6-expressing neurites were manually segmented, and pixel-wise ratiometric YFP/CFP images were generated. For each animal, multiple neurites were sampled per imaging session. A neurite was defined as having calcium overload if its mean YFP/CFP ratio exceeded 1.79, corresponding to intracellular calcium levels >235 nM. The proportion of overloaded neurites was calculated per mouse and used as the primary outcome measure of treatment efficacy.

### 2.5. Behavioral Testing

#### 2.5.1. Open Field Test

Mice were individually placed in a 27 × 27 × 20 cm acrylic arena and allowed to explore freely for 10 minutes. Locomotion was tracked via video recording and analyzed with EthoVision XT software (Noldus). Total distance traveled, time spent in the center zone, and center entry frequency were quantified to evaluate locomotor activity and anxiety-like behavior

#### 2.5.2. Y-Maze Test

Each mouse was introduced to the center of a Y-maze (arm dimensions: 5 × 30 × 12 cm; arms arranged at 120° angles) for a 10-minute exploration period. An arm entry required all four limbs to cross into an arm. Spontaneous alternation (SA) percentage was calculated as SA% = [Number of triads / (Total entries − 2)] × 100, with a triad defined as consecutive entries into three different arms. The maze was disinfected with 70% ethanol between tests.

#### 2.5.3. Fear Conditioning Test

Training took place in chambers (30 × 24 × 21 cm; MED-Associates) equipped with a stainless-steel grid floor and auditory stimulation system. Following a 2-minute habituation, mice received three tone-shock pairings (80 dB, 30 s tone; 0.6 mA shock during the final 2 s, co-terminating with the tone), separated by 210-second intervals. Training was conducted between 5:00 and 7:00 pm.

For contextual recall, mice were returned to the same chambers 24 hours later and observed for 3 minutes without tone or shock. One hour later, cued recall was assessed in a modified environment with altered floor and wall features. After 3 minutes of exploration, a 30-second tone was presented without shock. Freezing behavior was automatically analyzed using Video Freeze software (Med Associates Inc.). Chambers were sanitized with 70% ethanol between sessions.

### 2.6. Computational methods

#### 2.6.1. High-throughput virtual screening

For protein-ligand docking analysis, 3D structure of intact GABA_A_ receptor unit was retrieved from PDB databank (https://www.rcsb.org/) with accession ID 6D6U[23]. FDA-approved drug library was retrieved from DrugBank database (https://go.drugbank.com/). GABA_A_ receptor structure was prepared before docking analysis, using Protein Preparation Wizard from Schrodinger Maestro SBDD suite. Since the retrieved structure has a bound ligand (flumazenil), before screening, a short simulation for 100ns was conducted without any intact ligand, using Desmond simulation package to relax the protein complex. Retrieved FDA-approved drug library was prepared using LigPrep module from Schrodinger Maestro SBDD suite. For high-throughput virtual screening, Glide algorithm from Schrodinger Maestro SBDD suite was used[24]. Docking grid-box was generated based on the information from bound ligand (flumazenil) in the retrieved PDB structure (6D6U). The first stage docking was conducted using standard precision mode, followed by high-precision mode. For Gibbs binding free energy, MMGBSA method was used via Schrodinger Prime module[24].

#### 2.6.2. Machine learning and Neural network analysis

For predicting GABA_A_ receptor agonists and antagonists via machine learning approach, known GABA_A_ receptor agonists and antagonists were retrieved from PubChem database as smiles format and converted to SDF format using OpenBabel. Molecular descriptors (both 2D and 3D) for the retrieved compounds were calculated using QikProp module from Schrodinger SBDD suite[25]. Machine learning and neural network analysis were performed using Orange data mining software[26, 27].

#### 2.6.3. Molecular Dynamic Simulation

Molecular Dynamic (MD) simulations were performed using Desmond simulation package [28, 29]. Protein-ligand complex was first prepared using Protein Preparation Wizard from Maestro Visualizer. To mimic the physiological conditions, MD simulations of GABA_A_ receptor::drug complex were performed in a virtual lipid bilayer (cell membrane) system. GABA_A_ receptor::drug complex was placed in a POPC lipid bilayer system in such a way that the lipid bilayer covers the transmembrane region of GABA_A_ receptor. This GABA_A_ receptor::drug complex with intact lipid bilayer was then immersed in a triclinic simulation box containing Simple Point Charge (SPC) water. Simulation system was filled with 0.15M NaCl, excluding the counterions added to neutralize the system. After energy minimization (5000 steps) and equilibration (300ps), the whole system was simulated for 200ns. Simulation trajectories were analyzed for movement of chloride ions in and out of the membrane pore present at the M2 transmembrane region of GABA_A._

### 2.7. Statistical Analysis

Statistical analyses were performed using GraphPad Prism 9. Data were first tested for normality (Shapiro–Wilk test). Depending on distribution, comparisons were made using parametric (unpaired t-test, two-way ANOVA with Tukey’s) or non-parametric (Mann–Whitney U test, Kruskal–Wallis test followed by Dunn’s multiple comparisons test). Results are presented as mean ± SEM unless otherwise specified. P-values < 0.05 were considered statistically significant. All analyses were conducted with the experimenter blinded to treatment group.

## 3. RESULTS

### 3.1. Computational Modeling Identified Structural Dynamics of Ligand Binding to GABA_A_ Receptor

Young APP mice exhibit NREM sleep deficits and impairments of slow oscillation due to deficits in GABA and GABA receptor expression. Thus, we set out to perform an unbiased computational screen to identify an FDA-approved compound that could potentiate GABA_A_ signaling, rescue sleep and slow AD.

To investigate the structural dynamics involved in ligand binding to the GABA_A_ receptor, the three-dimensional quaternary structure of the receptor was retrieved from the Protein Data Bank (PDB). The structure comprises two α subunits, two β subunits, and one γ subunit. Structural visualization revealed the presence of a membrane pore (M2) and the benzodiazepine (BZD) binding site (Fig. 1A). The BZD binding site is located at the interface between the α and γ subunits of the GABA_A_ receptor (Fig. 1A). Since both GABA_A_ receptor agonists and antagonists primarily target the BZD binding region[23, 30], docking calculations were performed using known positive allosteric modulators (PAMs), e.g., diazepam, and competitive antagonists, e.g., flumazenil (FYP) as standard ligands targeting the BZD binding site. The term “agonist” here refers to PAMs that enhance GABA inhibitory action. The term “antagonist” refers to competitive antagonists, drugs that inhibit GABA inhibitory action. Docking results predicted binding affinities of -38.91 kcal/mol for diazepam and -27.56 kcal/mol for flumazenil at the BZD site (Supplementary Fig. 1A).

**Figure 1.**
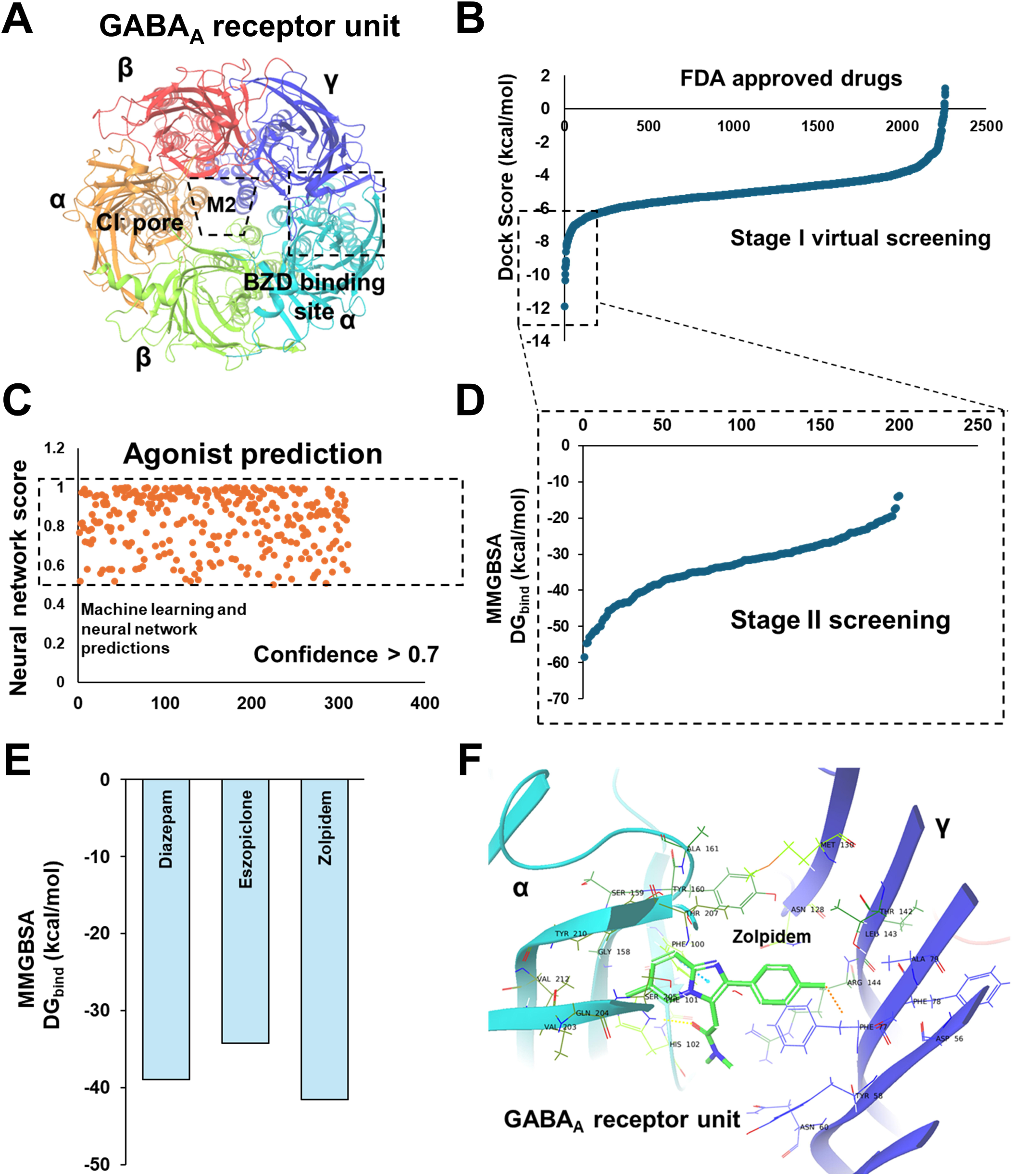
Quaternary structure and functional dynamics of the GABA_A_ receptor. **(A)** Experimental structure of the GABA_A_ receptor showing distinct subunits (colored based on GABA_A_ receptor subunits, BZD, α, β, and Υ), forming the quaternary assembly. The ligand-binding site, Benzodiazepine (BZD) binding site, is delineated with a box. **(B)** Predicted docking scores for the FDA-approved drug library, ranked from highest to lowest interaction energies during Stage I of the screen. **(C)** Machine learning and neural network-based classification of the top 300 compounds from (**B**), distinguishing predicted GABA_A_ receptor agonists from antagonists. (**D**) Predicted Molecular Mechanics-Generalized Born Solvent Accessible (MM-GBSA)-based Gibbs binding free energies for the top 200 compounds identified in (B) during Stage I of the screen. **(E)** Comparison of predicted Gibbs binding free energies for zolpidem and eszopiclone with that of diazepam. **(F)** Predicted binding pose of zolpidem in complex with the GABA_A_ receptor, highlighting interacting residues within the BZD binding site.

To further understand the dynamics of ligand binding, 200 ns atomistic molecular dynamics (MD) simulations were performed on the docked complexes of the GABA_A_ receptor bound to either diazepam or flumazenil, with the receptor embedded in a virtual lipid membrane (Supplementary Fig. 1B). Since the chloride transport through the M2 membrane pore is one of the key functional properties of the GABA_A_ receptor, the number of chloride ions passing through the M2 pore during the simulations was analyzed. Diazepam facilitated chloride ion transport across the membrane throughout the 200 ns simulation (Supplementary Fig. 1C). In contrast, flumazenil inhibited chloride ion transport across the membrane when bound to the GABA_A_ receptor (Supplementary Fig. 1D).

### 3.2. High-throughput Virtual Screening and Neural Network Predicted High-affinity GABA_A_ Receptor Modulators

To identify drug molecules targeting the GABA_A_ receptor as potentiators, we computationally screened an FDA-approved drug library against the BZD-binding site, based on the docking models as shown in Fig. 1B. The docking protocol was conducted in two stages. In Stage I, a high-throughput (standard-precision) docking approach was used to screen approximately 2,300 FDA-approved drugs against the GABA_A_ receptor. Drug candidates were ranked based on docking scores (Fig. 1B). Since both agonists and antagonists bind at the BZD-binding site, machine learning (ML) and neural network models were employed to classify potential agonists among the top-ranked compounds. For model training, we curated a dataset of 32 known GABA_A_ receptor agonists and 24 antagonists from public sources, including the PubChem database. Multiple ML algorithms were evaluated, including Support Vector Machine (SVM), Logistic Regression, Random Forest, and Neural Networks. Molecular descriptors (∼277) were computed for all compounds, both training and unknown, using the Mordred tool. Model development, training, and predictions were conducted using Orange Data Mining.

Model performance was assessed via random sampling. Among the tested models, SVM and Neural Networks showed the highest prediction accuracies, with SVM achieving 75.3% accuracy (AUC: 80.3%) and the Neural Network model achieving 74.4% accuracy (AUC: 77.3%). Other models, including Logistic Regression, showed accuracies below 70% and were therefore excluded from further analysis. Model training was repeated multiple times to ensure robustness, with SVM and Neural Network consistently demonstrating strong predictive performance. Once trained, these models were used to predict the agonist/antagonist classification for the full set of 2,256 FDA-approved therapeutics during Stage I. To minimize false positives, only compounds predicted as agonists by both the SVM and Neural Network models were selected, yielding 311 candidates (Fig. 1C).

To further refine the predictions, the top 200 compounds (top 10% from Stage I) were subjected to Stage II screening using the Molecular Mechanics Generalized Born Solvent (MM-GBSA) method. Drug candidates were ranked based on estimated Gibbs free energy of binding (Fig. 1D). Using predictions from both machine learning (properties) and computational docking (binding affinity) estimates (Fig. 1E), as well as its hypnotic properties, Zolpidem was selected as the leading drug candidate for further investigation.

### 3.3. MD Simulation Predicted Stability in Zolpidem’s Binding to GABA_A_ Receptor and Facilitating Chloride Ion Transport

Protein-ligand docking analysis of the GABA_A_ receptor bound to target drug molecules identified key amino acid residues involved in ligand recognition. Diazepam interacted with γPhe77, α1Thr162, α1Ser204, and α1Ser206 (Supplementary Fig. 2A-B). Protein-ligand interaction profiling of GABA_A_ receptor bound to zolpidem showed similar interaction pattern at the BZD-binding site when compared to diazepam (Fig. 1F).

To evaluate the binding stability of zolpidem and its potential to facilitate chloride ion transport across the membrane, a 200 ns molecular dynamics (MD) simulation of the GABA_A_ receptor– zolpidem complex was performed, with the receptor embedded in a virtual lipid bilayer. Simulation trajectories were analyzed to assess binding stability and the number of contacts between chloride ions and the M2 transmembrane region (chloride pore) of the receptor. Zolpidem maintained stable binding at the BZD site throughout the 200 ns simulation (Supplementary Fig. 3A).

Snapshots from simulation trajectories at different time points revealed that zolpidem binding induced structural changes in the GABA_A_ receptor, notably the formation of a membrane pore (Supplementary Fig. 3B). In contrast, competitive antagonist flumazenil inhibited such membrane pore formation during the 200 ns simulation (Supplementary Fig. 3C).

Since zolpidem induced a membrane pore formation, we next examined whether it could facilitate chloride ion transport. Analysis of the simulation trajectories for the number of chloride ions passing through M2 membrane showed that zolpidem facilitated chloride ion transport across the membrane, with an increased number of chloride ions passing through the M2 membrane pore over the course of the 200 ns simulation (Supplementary Fig. 4A, B). Visualization of chloride ion trajectories indicated Cl^-^ ions traversed the virtual membrane from the cytoplasmic side to the extracellular side, interacting with M2 transmembrane residues (Supplementary Fig. 4A). Thus, as a result of unbiased computational screen, zolpidem emerged as a leading candidate potentiating GABA_A_ signaling. Zolpidem’s effect on slow oscillation, sleep and Alzheimer’s progression was tested in a mouse model of amyloidosis, young APP mice.

### 3.4. Zolpidem Restored Slow Oscillation in APP Mice

APP mice exhibit slow oscillation of low power compared to that of nontransgenic littermate controls [19]. To determine the extent to which zolpidem restores slow oscillation in APP mice, we first recorded baseline cortical slow wave activity using voltage-sensitive dye (VSD) through cranial windows via wide-field microscopy in vivo. Subsequently, zolpidem at three different concentrations (3, 10, 30 mg/kg) or vehicle control (0 mg/kg) was administered intraperitoneally (i.p.) to APP mice (Fig. 2A).

**Figure 2.**
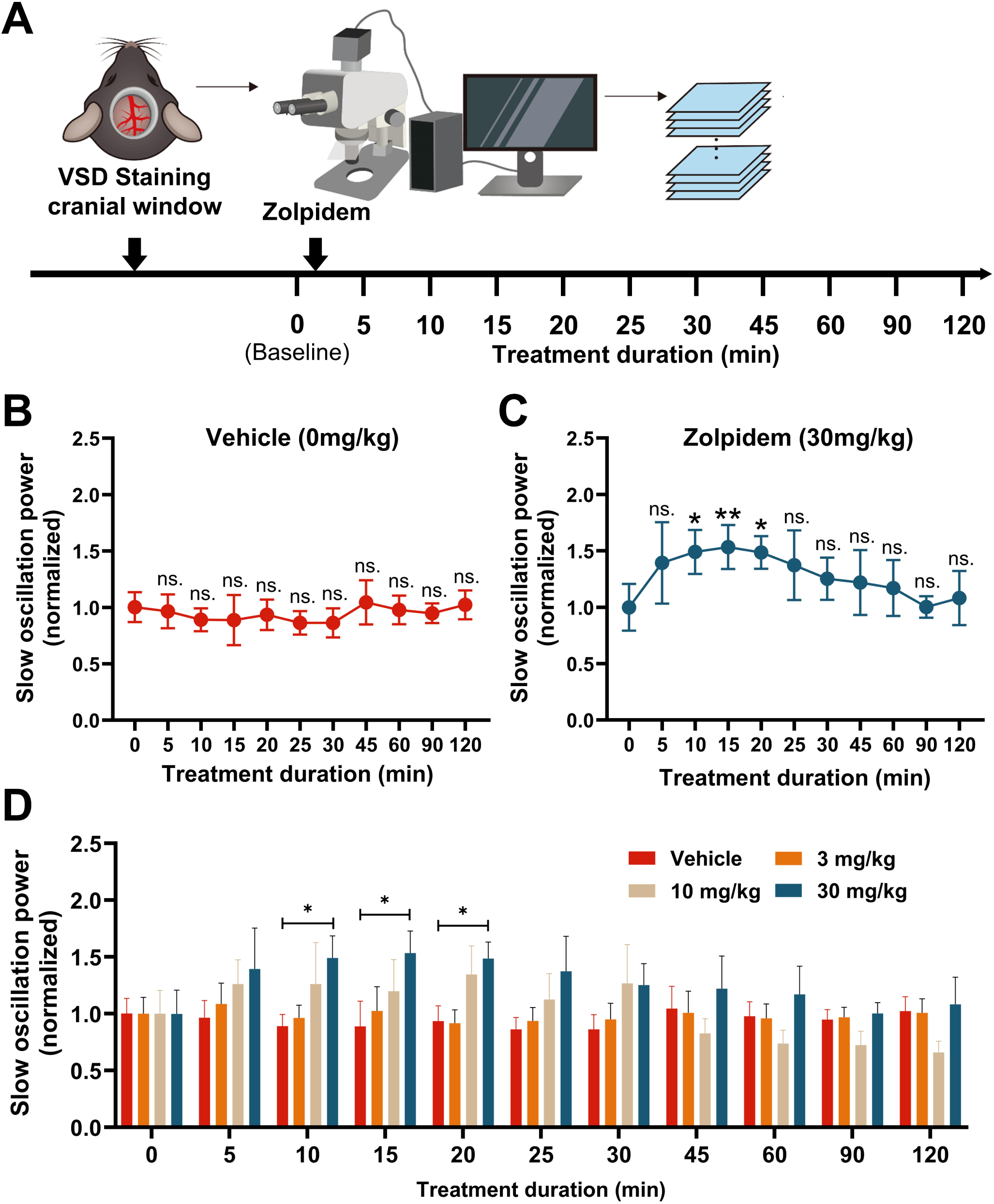
Voltage-sensitive dye (VSD) imaging revealed zolpidem-induced enhancement of cortical slow oscillation. **(A)** Schematic of VSD imaging setup over somatosensory cortex in an APP mouse. **(B)** Statistical comparison of slow oscillation (0.5–1 Hz) power before and after vehicle (0 mg/kg) treatment. Friedman test followed by Dunn’s multiple comparisons test. **(C)** Statistical comparison of slow oscillation (0.5–1 Hz) power before and after 30 mg/kg zolpidem treatment. Friedman test followed by Dunn’s multiple comparisons test. **(D)** Time course of normalized slow oscillation power across 120 minutes post-gavage in vehicle (0 mg/kg), 10 mg/kg, and 30 mg/kg treated groups. one-way ANOVA followed by Tukey’s post hoc multiple comparisons test. Data shown as mean ± SEM (n=5 mice/group). *p < 0.05, **p < 0.01. n.s., non-significant. Lack of stars indicates lack of significance.

Given the rapid permeability of zolpidem across the blood-brain barrier [31, 32], recordings were taken at 5-minute intervals within the first 30 minutes post-injection, and at longer intervals up to 120 minutes post-injection. We observed a gradual increase in slow oscillation power after zolpidem administration, reaching a significant elevation relative to baseline at 10, 15, and 20 minutes with a 30 mg/kg dose. Subsequently, slow oscillation power gradually declined, returning to baseline levels (Fig. 2C). Meanwhile, APP mice treated with vehicle control showed no significant changes in slow oscillation power throughout the 120-minute recording period following injection (Fig. 2B). When directly compared with vehicle-treated controls, APP mice treated with 30 mg/kg zolpidem exhibited significantly enhanced slow oscillation power at 10, 15, and 20 minutes post-injection. However, lower concentrations of zolpidem (3 and 10 mg/kg) did not produce significant changes in slow oscillation power compared to baseline or the vehicle group (Supplemental Fig. 5). Therefore, we selected 30 mg/kg zolpidem for subsequent experiments.

### 3.5. Zolpidem Ameliorated NREM Sleep Deficits in APP Mice

APP mice spend less time in NREM sleep and more time Awake compared to nontransgenic littermate controls at 6 months of age [19]. To determine whether zolpidem could rescue NREM sleep deficits in APP mice, we surgically implanted wireless EEG/EMG probes and performed recordings to monitor sleep architecture in freely moving animals (Fig. 3A). 3-6 month old APP mice were randomly assigned into one of two groups: vehicle-treated controls or mice receiving 30 mg/kg zolpidem consecutively for 2 days. Baseline sleep patterns were recorded on Day 0 prior to intervention. On Day 1 and Day 2, animals received one injection per day at around ZT2 (2 hours after lights-on, corresponding to 9 AM) (Fig. 3B, C). EEG/EMG recordings were taken for 2 days post-injection, on Day 3 and Day 4, to monitor the lasting effects of the drug.

**Figure 3.**
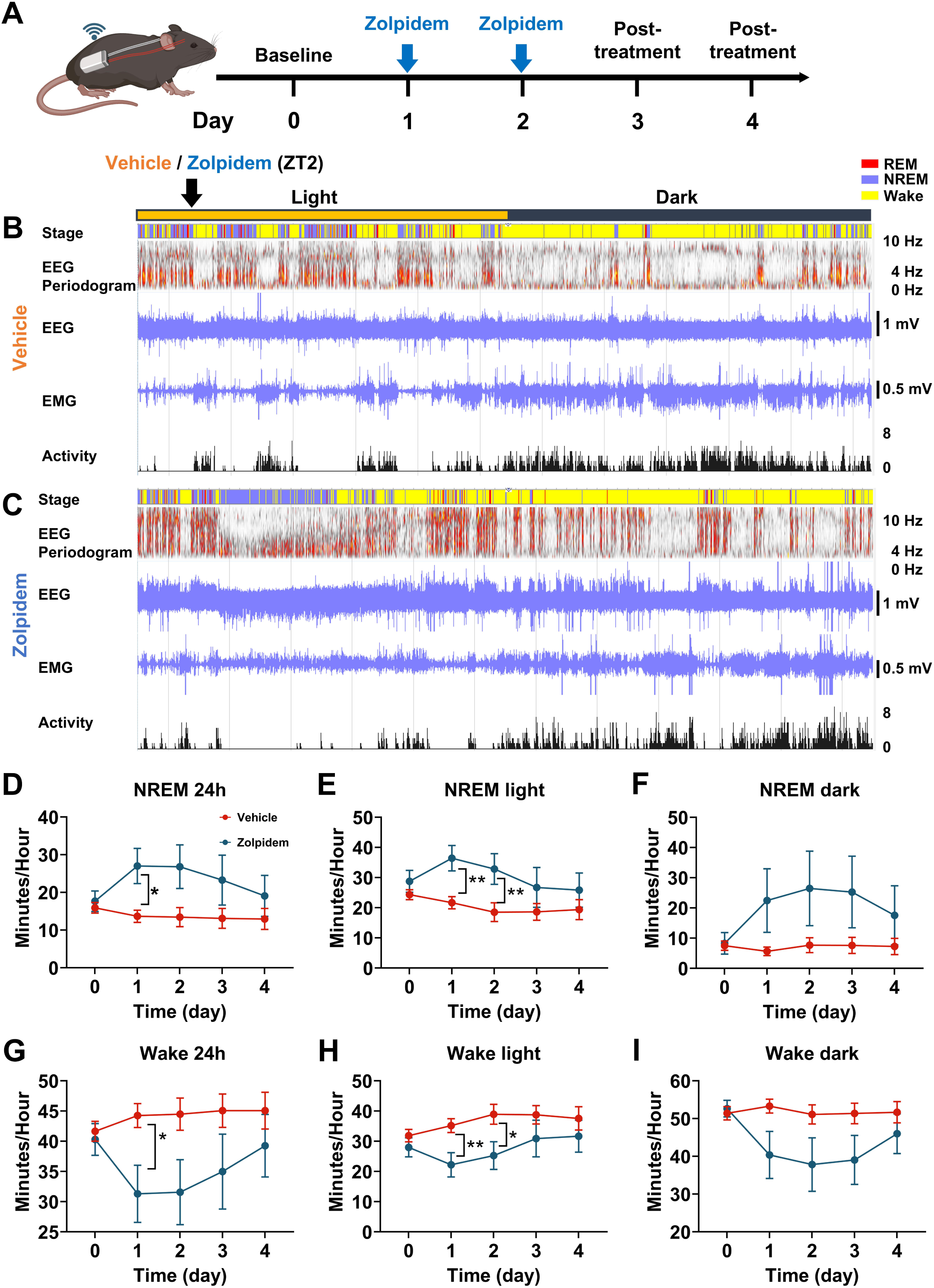
Zolpidem acutely increased NREM sleep and reduced wakefulness in APP mice. **(A)** Experimental timeline for EEG/EMG recordings showing baseline, zolpidem administration on days 1 and 2, and post-treatment recording on days 3 and 4. **(B)** Representative hypnogram, EEG spectrogram, EEG/EMG traces, and locomotor activity for a vehicle-treated mouse. **(C)** Corresponding plots for a zolpidem-treated mouse, demonstrating increased NREM sleep (blue) and reduced wakefulness (yellow) during the light phase. **(D-F)** Quantification of NREM sleep durations across 24 hours (**D**), light phase (**E**), and dark phase (**F**). **(G-I)** Quantification of wakefulness durations across 24 hours (**G**), light phase (**H**), and dark phase (**I**). Mixed-effects analysis followed by Tukey’s multiple comparisons test. Black* indicates significant group differences between zolpidem and vehicle treatments. Data shown as mean ± SEM (n=6-7 mice/group). *p < 0.05; **p < 0.01. Lack of stars indicates lack of significance.

Compared to vehicle-treated APP mice, zolpidem treatment significantly increased NREM sleep duration and reduced wakefulness during the 24 hours (Fig. 3D, G). That effect was prevalent during the light phase, when mice spent more time for NREM sleep (Fig. 3E, H). However, the effect of zolpidem was absent during the dark phase when animals were more active (Fig. 3F, I). This suggested short-lived effects of the drug and limited toxicity. Drug effects lasted through the light phase and did not carry over into the active dark phase. The sleep-inducing effects of zolpidem disappeared as early as Day 3, the day after the final injection, suggesting a short-lasting impact of zolpidem on sleep architecture. No significant group differences were observed in REM sleep across all time points (Supplemental Fig. 6A–C). These findings indicate that zolpidem can restore NREM sleep deficits in APP mice and not interfere with the active periods.

### 3.6. Zolpidem Enhanced Slow Oscillation and Dynamically Modulated High-Frequency Rhythms in APP Mice

Our published findings demonstrate that young APP mice exhibit reduced slow wave power (0.5-1Hz) and elevated higher-frequency components during sleep, indicative of cortical hyperexcitability, an aberration commonly observed in young amyloidosis models[19]. To further investigate the spectral dynamics following zolpidem administration, we analyzed EEG power within the first 6 hours after the drug was administered. This window captures the immediate pharmacological effects of zolpidem during the period when mice are naturally more likely to sleep. We performed Fourier transform analysis on EEG data acquired within the first 6 hours post-injection. This allowed us to determine the drug effects on cortical oscillatory activity. As shown in Fig. 4A and B, compared to their own baseline, zolpidem treatment significantly increased slow oscillation (0.5-1 Hz) and delta (1-4 Hz) power during the first 2-3 hours post-injection. This reflected enhanced low-frequency activity associated with deep NREM sleep. Vehicle-treated mice showed no significant change from baseline at any time point. Moreover, direct comparisons between groups revealed that the slow oscillation power was significantly higher in the zolpidem group than in the vehicle group at 2 hours post-injection (Fig. 4A).

**Figure 4.**
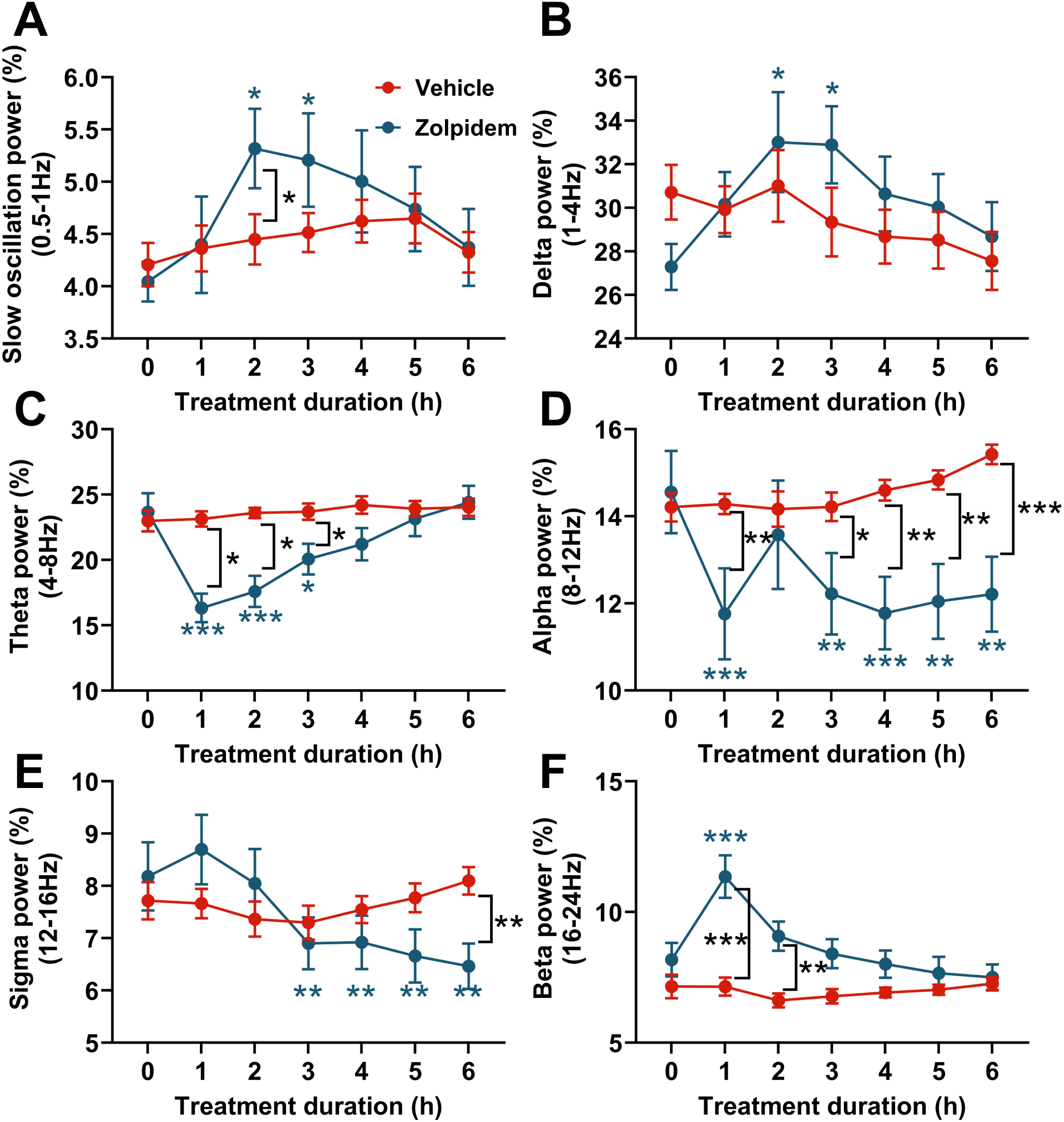
Zolpidem dynamically modulated cortical oscillatory activity across multiple frequency bands in APP mice during NREM sleep. We analyzed EEG power during NREM sleep within the first 6 hours after the drug was administered and presented the hourly changes. (A) Slow oscillation power (0.5–1 Hz) across treatment groups. (B) Delta power (1–4 Hz) across treatment groups. (C) Theta power (4–8 Hz) across treatment groups. (D) Alpha power (8–12 Hz) across treatment groups. (E) Sigma power (12–16 Hz) across treatment groups. (F) Beta power (16–24 Hz) across treatment groups. Two-way ANOVA followed by Tukey’s multiple comparisons test. Blue* indicates significant differences compared to baseline within the zolpidem group. Black* indicates significant group differences between zolpidem and vehicle treatments. Data shown as mean ± SEM (n=6-7 mice/group). *p < 0.05; **p < 0.01; ***p < 0.001. Lack of stars indicates lack of significance.

For higher-frequency bands, zolpidem significantly suppressed theta (4–8 Hz) during 1-3 hours post-injection compared to baseline (Fig. 4C). Alpha (8–12 Hz) was suppressed 1, 3-6 hours post-injection compared to baseline (Fig. 4D). Sigma (12–16 Hz) activity was suppressed 3-6 hours post-injection relative to baseline (Fig. 4E). The vehicle group showed no significant changes from baseline in these bands. Importantly, zolpidem-treated mice exhibited significantly lower power than vehicle-treated mice at hours 1-3 for theta, hours 1, 3-6 for alpha, hour 6 for sigma (p < 0.05) (Fig. 4C-D). Thus, zolpidem facilitated lower brain rhythms and effectively suppressed higher rhythms. In contrast, beta (16–24 Hz) power transiently increased in the zolpidem group during the first hour, relative to both baseline and vehicle controls (p < 0.05), but returned to baseline thereafter (Fig. 4F). Together, these results indicate that zolpidem alters the spectral profile in APP mice by promoting slow oscillatory activity while dynamically modulating higher-frequency rhythms in a time-dependent manner.

### 3.7. Chronic Zolpidem Treatment Suppressed Amyloid Deposition in APP Mice

Building on the sleep-enhancing effects of acute zolpidem treatment, we next sought to examine whether sustained treatment could affect Alzheimer’s disease pathology. Given the causal link between NREM sleep impairments and accelerated Aβ deposition[33], we assessed the impact of chronic daily zolpidem administration at 30mg/kg on amyloid burden at baseline, Weeks 2 and 4 of a 4-week treatment period (Fig. 5A).

**Figure 5.**
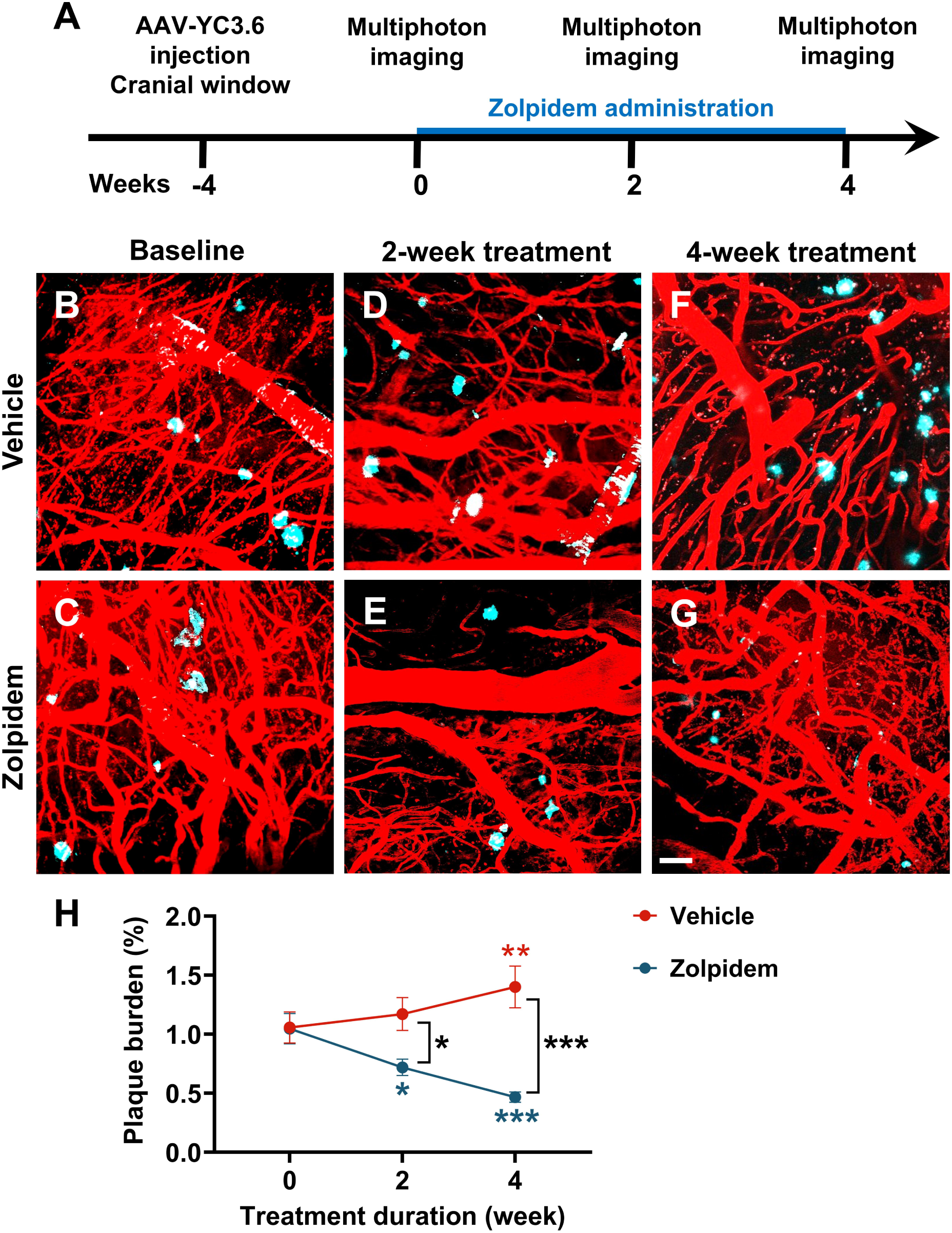
Chronic zolpidem treatment reduced amyloid plaque burden in APP mice. **(A)** Experimental timeline of multiphoton imaging at baseline, 2 weeks, and 4 weeks of treatment. **(B-G)** Representative two-photon images of amyloid plaques (cyan, Methoxy-X04 labeling) and cortical vasculature (red, Texas Red dextran) at baseline (**B, C**), after 2 weeks of treatment (**D, E**), and after 4 weeks of treatment (**F, G**) for vehicle (**B, D, F**) and zolpidem-treated mice (**C, E, G**). Zolpidem treatment visibly reduced plaque accumulation over time. **(H)** Quantification of cortical plaque burden over the 4-week treatment period. Zolpidem significantly reduced amyloid plaque burden compared to both baseline and vehicle-treated controls. Mixed-effects analysis followed by Tukey’s multiple comparisons test. Blue* indicates significant differences compared to baseline within the zolpidem group. Red* indicates significant differences compared to baseline within the vehicle group. Black* indicates significant group differences between zolpidem and vehicle treatments. Data shown as mean ± SEM (n=6-7 mice/group). *p < 0.05; **p < 0.01; ***p < 0.001.

To visualize amyloid plaques in vivo over time, amyloid deposits were labeled with Methoxy-X04. Plaques were visualized in anesthetized mice through cranial windows using multiphoton microscopy. Baseline imaging (Fig. 5B, C) was performed before the start of treatment, followed by repeated imaging of same animals at Weeks 2 (Fig. 5D, E) and 4 (Fig. 5F, G). Plaque burden was comparable between groups at baseline. In vehicle-treated APP mice, plaque burden gradually increased, with a significant elevation at Week 4 compared to baseline. In contrast, zolpidem-treated mice exhibited a substantial reduction in plaque burden beginning at Week 2, which continued to decline through Week 4. A significant difference in plaque burden between zolpidem-and vehicle-treated mice was observed at Week 2 and became more pronounced by Week 4 (Fig. 5H). Thus, chronic zolpidem treatment effectively suppressed Aβ accumulations.

### 3.8. Chronic Zolpidem Treatment Reduced the Percentage of Neuronal Processes with Calcium Overload in APP Mice

Calcium homeostasis is necessary for proper neuronal function. Neuronal calcium homeostasis is impaired in APP mice. A subset of cortical neurons exhibits calcium overload, or significant elevations in basal calcium, due to amyloid β-induced dysregulation, particularly within neurites, or neuronal processes of APP mice [34, 35]. To evaluate whether chronic zolpidem treatment can restore neuronal calcium homeostasis, we expressed the ratiometric calcium indicator Yellow Cameleon 3.6 (YC3.6) in the cortex of APP mice. YC3.6 is a genetically encoded FRET-based sensor containing YFP and CFP fluorophores. YC3.6 allows determination of real-time intracellular calcium levels. Neurites were classified as overloaded when the YFP/CFP ratio exceeded 1.79, corresponding to intracellular calcium concentrations greater than 235 nM, or two standard deviations above the mean of neurites in nontransgenic controls [22].

Representative images show numerous, red-labeled neurites with calcium overload (yellow arrows) in vehicle-treated mice at 2 and 4 weeks (Fig. 6A, C, E). However, zolpidem-treated mice showed a marked reduction in overloaded neurites over the same time period (Fig. 6B, D, F). Histograms of YFP/CFP ratio distributions showed a progressive increase in the percentage of calcium-overloaded neurites in the vehicle-treated group over time (Fig. 6G, red box). However, zolpidem-treated mice exhibited a gradual reduction in the percentage if overloaded neurites over the treatment period (Fig. 6H, red box). Quantitative analysis demonstrated that the percentage of neurites exhibiting calcium overload increased in vehicle-treated mice from baseline to Week 2 and increased further at Week 4. However, zolpidem treatment significantly reduced the percentage of neurites with calcium overload by Week 4, compared to baseline and vehicle controls (Fig. 6I). These findings suggest that chronic zolpidem treatment restored calcium homeostasis and prevented calcium overload in neurites of APP mice, providing evidence for its potential neuroprotective effects beyond sleep regulation.

**Figure 6.**
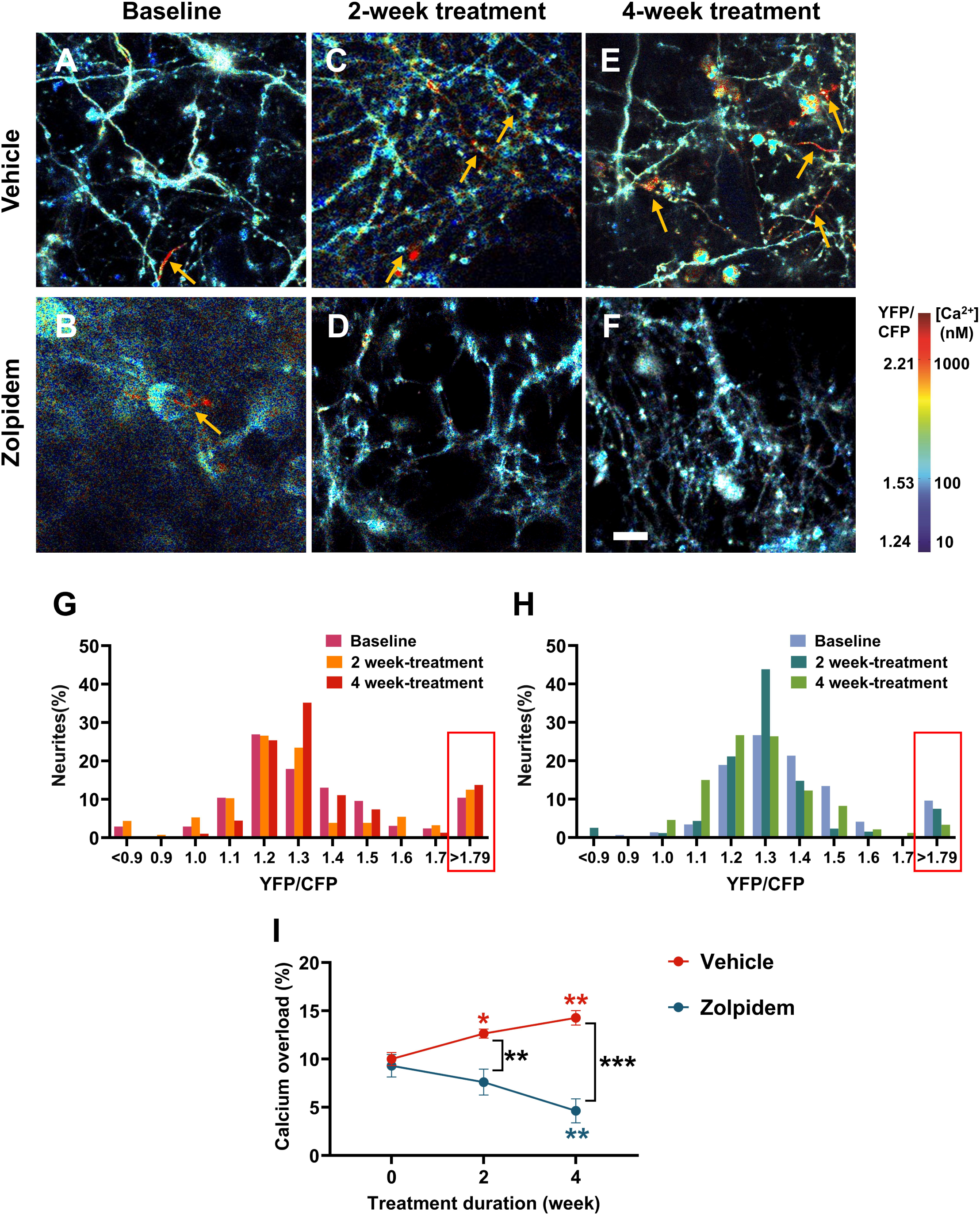
Zolpidem restored neuronal calcium homeostasis in APP mice. **(A–F)** Representative images of YC3.6-labeled neurites showing calcium overload (red) in vehicle and zolpidem groups at baseline, 2 weeks, and 4 weeks. **(G–H)** Histograms of YFP/CFP ratios within neurite populations treated with vehicle (**G**) and zolpidem (**H**). **(I)** Quantification of calcium-overloaded neurites (%) across groups. Mixed-effects analysis followed by Tukey’s multiple comparisons test. Blue* indicates significant differences compared to baseline within the zolpidem group. Red* indicates significant differences between the zolpidem group and vehicle group at the indicated time point. Black* indicates significant group differences between zolpidem and vehicle treatments. Data shown as mean ± SEM (n=6-7 mice/group). *p < 0.05; **p < 0.01; ***p < 0.001.

### 3.9. Chronic Zolpidem Treatment Improved Memory Consolidation in APP Mice

To determine whether chronic zolpidem treatment alters cognitive function in APP mice, we conducted a series of behavioral assays, including open field, Y-maze, and fear conditioning tests, following 4 weeks of daily treatment with either 30 mg/kg zolpidem or vehicle. In the open field test (Fig. 7A), zolpidem- and vehicle-treated mice exhibited comparable total distance traveled, average velocity, and frequency of center entries (Fig. 7B–D), indicating no significant differences in locomotor activity or anxiety-like behavior between groups. Similarly, no significant differences were observed in spontaneous alternations in the Y-maze test (Fig. 7E, F), suggesting that zolpidem treatment did not impact working memory in APP mice under these conditions.

**Figure 7.**
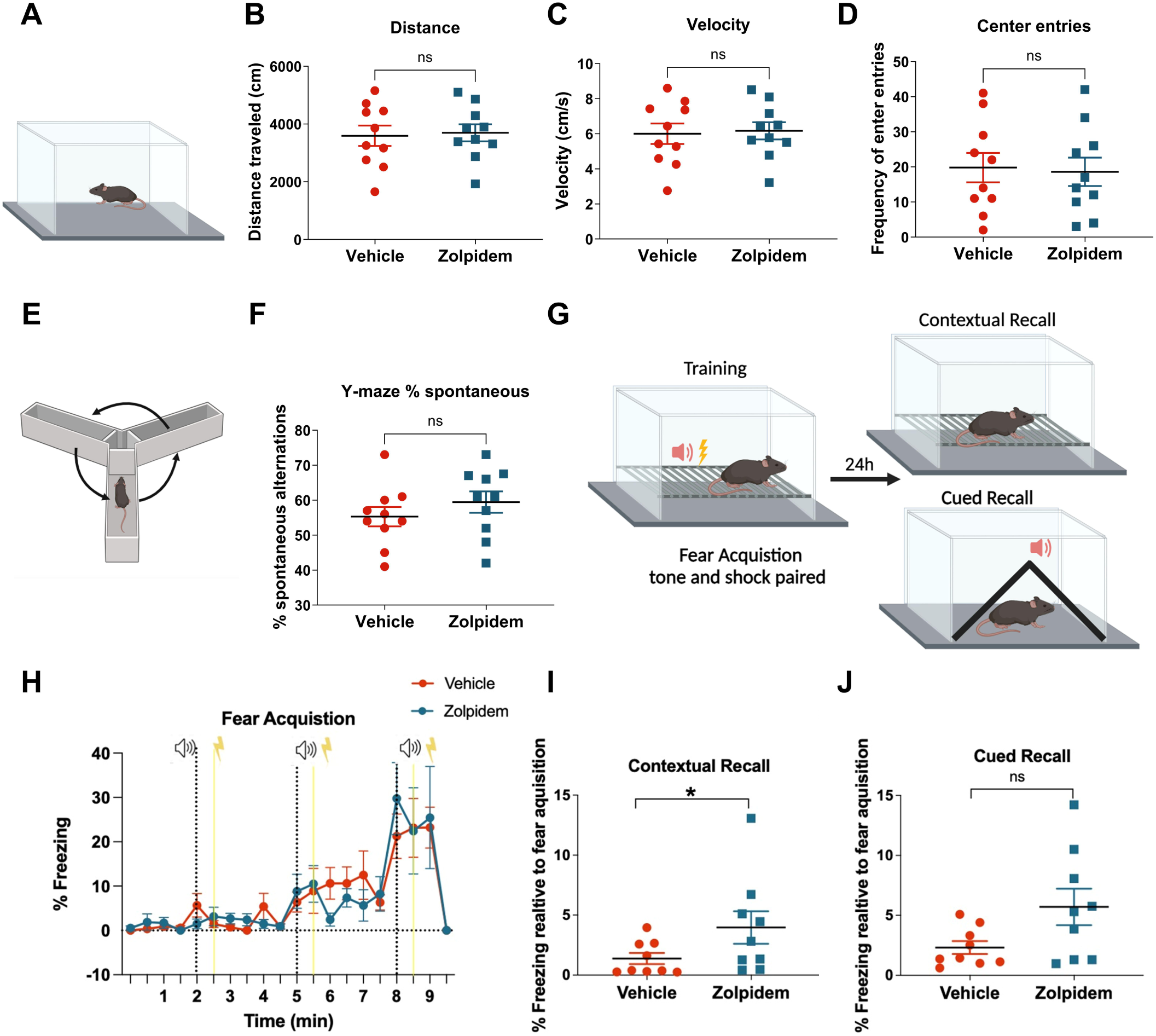
Zolpidem improved contextual memory without affecting locomotor or working memory. **(A)** Experimental setup for open field. **(B–D)** Distance travelled (**B**), velocity (**C**) and center entries (**D**) as a measure of open field performance across groups. (**E**) Experimental setup for Y-maze test. **(F)** Y-maze alternation rates across groups. **(G)** Contextual fear conditioning setup. **(H-J)** Fear acquisition (**H**), contextual recall (**I**) and cued recall (**J**) across groups. Unpaired two-tailed t-test. Data shown as mean ± SEM (n=9-10 mice/group). *p < 0.05.

We next assessed sleep-dependent memory consolidation using a classical fear conditioning paradigm (Fig. 7G). During the acquisition phase, both groups exhibited increased freezing behavior over time (Fig. 7H), indicating successful formation of fear memory. Then animals were placed back into their home cages and allowed to consolidate fear memory during sleep. Twenty-four hours later, contextual fear recall was tested by reintroducing mice to the original training environment without any tone or shock. Zolpidem-treated mice demonstrated significantly higher freezing levels during contextual recall compared to vehicle-treated mice (Fig. 7I), suggesting improved memory consolidation. In contrast, no significant differences were observed between groups during cued recall in a novel context (Fig. 7J). These results suggest that chronic zolpidem administration selectively enhances contextual memory consolidation in APP mice without affecting general locomotion or working memory performance.

## 4. DISCUSSION

The current work identified that therapeutic targeting of sleep can profoundly impact the course of Alzheimer’s disease (AD) progression. We evaluated whether pharmacologically enhancing sleep-dependent brain rhythms using FDA-approved hypnotic zolpidem could restore cortical slow oscillation, improve NREM sleep duration, and alleviate key molecular and functional deficits in APP mice. Acute zolpidem administration markedly boosted slow oscillation power and enhanced sleep continuity, while chronic daily treatment over four weeks reduced amyloid plaque burden, normalization of neuronal calcium homeostasis, and improvements in memory consolidation without adversely affecting locomotor or general cognitive functions. These results underscore the therapeutic potential of pharmacologically restoring sleep-dependent brain rhythms to slow AD progression.

NREM sleep plays a key role in memory consolidation and the progression of AD [7]. Early-stage AD patients exhibit significant disruptions in NREM sleep, reduced slow-wave sleep, increased sleep fragmentation, and altered sleep architecture, linked to accelerated cognitive decline and worsening neuropathology [9, 36–38]. Among the core electrophysiological features of deep NREM sleep, slow oscillation (0.5-1 Hz) is a fundamental sleep-dependent brain rhythm. These large-amplitude, low-frequency cortical waves reflect highly synchronized neuronal activity, consisting of alternating down-states of neuronal silence and up-states of synchronous firing [10–12]. Slow oscillation is essential for memory consolidation [39], synaptic homeostasis [40], and metabolic waste clearance [41]. Impairments appear early in AD, often before substantial plaque deposition yet in the presence of oligomeric Aβ, suggesting a causal role in disease progression [42–44]. Our published work demonstrated that slow oscillation is impaired in APP mice, with cortical desynchronization, neuronal calcium overload, and accelerated amyloid pathology. Optogenetic restoration of slow oscillation slowed the AD progression, whereas further disruption worsened disease in APP mice [17, 18], indicating slow oscillation is a viable therapeutic target.

Deficits in inhibitory tone also contribute to slow wave disruptions during prodromal and early AD [19]. Exogenous GABA restored slow oscillation in an AD mouse model, whereas pharmacological blockade of GABA_A_ and GABAB receptors disrupted slow oscillation in healthy nontransgenic mice. Altered cortical inhibitory interneuron activity in APP mice further links interneuron dysfunction to impaired rhythms [17]. Using an optogenetic, we reported that rhythmic stimulation of GABAergic interneurons at the endogenous frequency of slow oscillation, 0.6 Hz, was sufficient to restore slow waves and sleep in APP mice [19]. Most importantly it slowed Alzheimer’s progression in APP mice. While this methodological strategy established a causal relationship between interneuron-dependent impairments in cortical slow wave activity and Alzheimer’s progression, its translational potential remained limited.

We therefore performed an unbiased computational screen to identify an FDA-approved compound that could rescue sleep, potentiate sleep-dependent brain rhythms and slow Alzheimer’s progression. Computational modeling revealed zolpidem’s strong and stable binding to the GABA_A_ receptor at the BZD site, with structural dynamics similar to diazepam. Molecular dynamics simulations showed that zolpidem facilitated chloride ion transport across the membrane, suggesting its ability to restore inhibitory tone. High-throughput docking and machine learning screening of FDA-approved drugs further prioritized zolpidem based on predicted binding affinity and potentiation of GABA signaling. These findings provided a mechanistic basis for testing zolpidem’s potential to rescue disrupted network activity in AD by enhancing GABAergic signaling and restoring cortical slow oscillation. To enhance translational impact based on these computational findings, we tested zolpidem, an FDA-approved GABA_A_ receptor modulator. Zolpidem is a proven hypnotic known to potentiate inhibition, promote sleep onset and maintain sleep clinically [45, 46]. Beyond its hypnotic effects, emerging evidence suggests that zolpidem has demonstrated beneficial neuromodulatory effects in various neurological disorders, including stroke [47, 48], traumatic brain injury [49, 50], and Parkinson’s disease [51, 52]. Although zolpidem has been shown to improve sleep quality in Alzheimer’s patients, its impact on slow oscillation dynamics and related AD pathology has not been thoroughly tested. As an initial step, we employed in vivo VSD imaging to directly assess the impact of different doses of zolpidem on cortical slow oscillation dynamics in APP mice. We identified 30 mg/kg as the most effective dose for enhancing slow oscillation power, which guided the design of our subsequent chronic treatment experiments.

Having established an effective dose for enhancing cortical slow oscillation, we next investigated whether zolpidem could alleviate the characteristic sleep disturbances observed in APP mice. Prior studies have shown that APP mice display fragmented sleep architecture, with increased wakefulness and reduced NREM sleep, particularly during the light phase, their typical rest period mirroring the sleep abnormalities seen in Alzheimer’s patients [19, 53, 54]. Using EEG/EMG recordings, we found that acute administration of 30 mg/kg zolpidem significantly increased NREM sleep during the 24-hour recording period, specifically during the light phase. These effects were not observed in vehicle-treated mice. Zolpidem effects dissipated by the following day. In addition, zolpidem had little to no impact on sleep architecture during the dark phase, the animals’ natural period of wakefulness, indicating that zolpidem selectively enhanced rest-phase sleep without disrupting activity during the dark phase.

Fourier transform analyses of EEG recordings revealed dynamic spectral changes induced by zolpidem treatment. Consistent with our previous findings of cortical hyperexcitability characterized by reduced slow-wave power and elevated higher-frequency rhythms in APP mice, zolpidem significantly increased slow oscillation (0.5–1 Hz) and delta (1–4 Hz) power within the initial hours post-injection. Conversely, zolpidem markedly suppressed higher-frequency rhythms, including theta (4–8 Hz), alpha (8–12 Hz), and sigma (12–16 Hz), while transiently elevating beta (16–24 Hz) activity in the first hour following administration. These spectral changes underscore zolpidem’s capacity to restore cortical excitatory-inhibitory balance, promoting deep sleep-associated low-frequency rhythms and dynamically modulating aberrant high-frequency activities prevalent in AD. These findings align with prior studies showing that zolpidem enhances slow-wave power and suppresses high-frequency EEG components during NREM sleep. For instance, previous work has demonstrated that zolpidem increases slow-wave (0.5-4 Hz) amplitude and power in rodents [55–57], corroborating the spectral changes observed in our APP mice. However, not all studies report consistent outcomes, some have observed reductions in SWA, likely due to differences in disease models, zolpidem dose, or the timing and duration of EEG analysis [58, 59].

To determine whether these acute effects could translate into longer-term benefits, we implemented a chronic dosing regimen in which APP mice received daily zolpidem injections for four weeks. Longitudinal two-photon imaging of amyloid plaques demonstrated that chronic zolpidem administration significantly suppressed amyloid deposition relative to vehicle-treated controls. This finding aligns with published studies indicating that impaired sleep exacerbates amyloid accumulation, whereas enhancing deep NREM sleep facilitates Aβ clearance [37, 60, 61].

In parallel, multiphoton calcium imaging revealed that chronic zolpidem treatment significantly reduced the proportion of neurites exhibiting calcium overload, a marker of synaptic dysfunction and neurotoxicity in AD [34, 35]. The observed decrease in the YFP/CFP ratios within neurites among zolpidem-treated mice suggests improved intracellular calcium homeostasis, which is typically disrupted by elevated amyloid-beta levels and associated cortical hyperexcitability [35]. Importantly, the normalization of neuronal calcium levels was evident as early as two weeks into treatment and became even more pronounced by week four, indicating a progressive, cumulative therapeutic effect.

Finally, to determine whether these neurophysiological improvements translated into functional gains, we conducted a battery of behavioral assays. Zolpidem-treated APP mice displayed intact locomotor function and healthy working memory, as indicated by open field and Y-maze performance. Notably, these mice exhibited enhanced contextual fear memory consolidation, suggesting that improved sleep and cortical network synchronization directly enhanced memory consolidation.

Zolpidem is FDA-approved hypnotic known to potentiate inhibitory tone and effective at inducing sleep in humans. Although previous studies have reported potential adverse effects of zolpidem treatment, including cognitive impairment and increased risk of falls in older adults, these outcomes appear to be dose-dependent and population-specific [62–64]. That is, these adverse outcomes could be attributed to long-treatment durations at high doses that induce toxicity. Indeed, zolpidem is routinely prescribed to elderly populations and has demonstrated a favorable safety profile in short-term intervention studies[63]. Supporting this, a recent randomized, triple-blind, placebo-controlled clinical trial found that zolpidem effectively decreased wakefulness and increased sleep duration in Alzheimer’s patients, with mild to no adverse events reported [65]. Other clinical and preclinical studies have similarly reported beneficial effects of chronic zolpidem administration. For example, long-term treatment with zolpidem has been shown to improve neurological function and promote behavioral recovery in stroke [47, 66]. In line with these findings, we did not observe any notable adverse effects on locomotion in our study. Short-duration treatment of 4 weeks at relatively low doses of 30mg/kg failed to elicit toxicity and adverse side-effects in APP mice as reported in present study. This may be attributable to factors such as differences in animal models, disease stage, dosing regimen, or treatment duration. Thus, intermittent low-dose administration at early AD stages, when cortical circuits are hyperactive from reduced inhibitory tone, may maximize benefit while minimizing sedation risk. Collectively, these observations highlight the importance of optimizing therapeutic strategies to maximize clinical benefit while minimizing potential risks.

## ACKNOWLEDGEMENT

The works was supported by is made to the donors of Alzheimer’s Disease Research, a program of BrightFocus Foundation, for support of this research (KVK, QZ).

## CONSENT STATEMENT

No human data was derived and, therefore, consent was not necessary.

