## Supplementary figures and images for "Zolpidem restores sleep and slows Alzheimer’s progression in a mouse model"

### Supplemental Figure 1

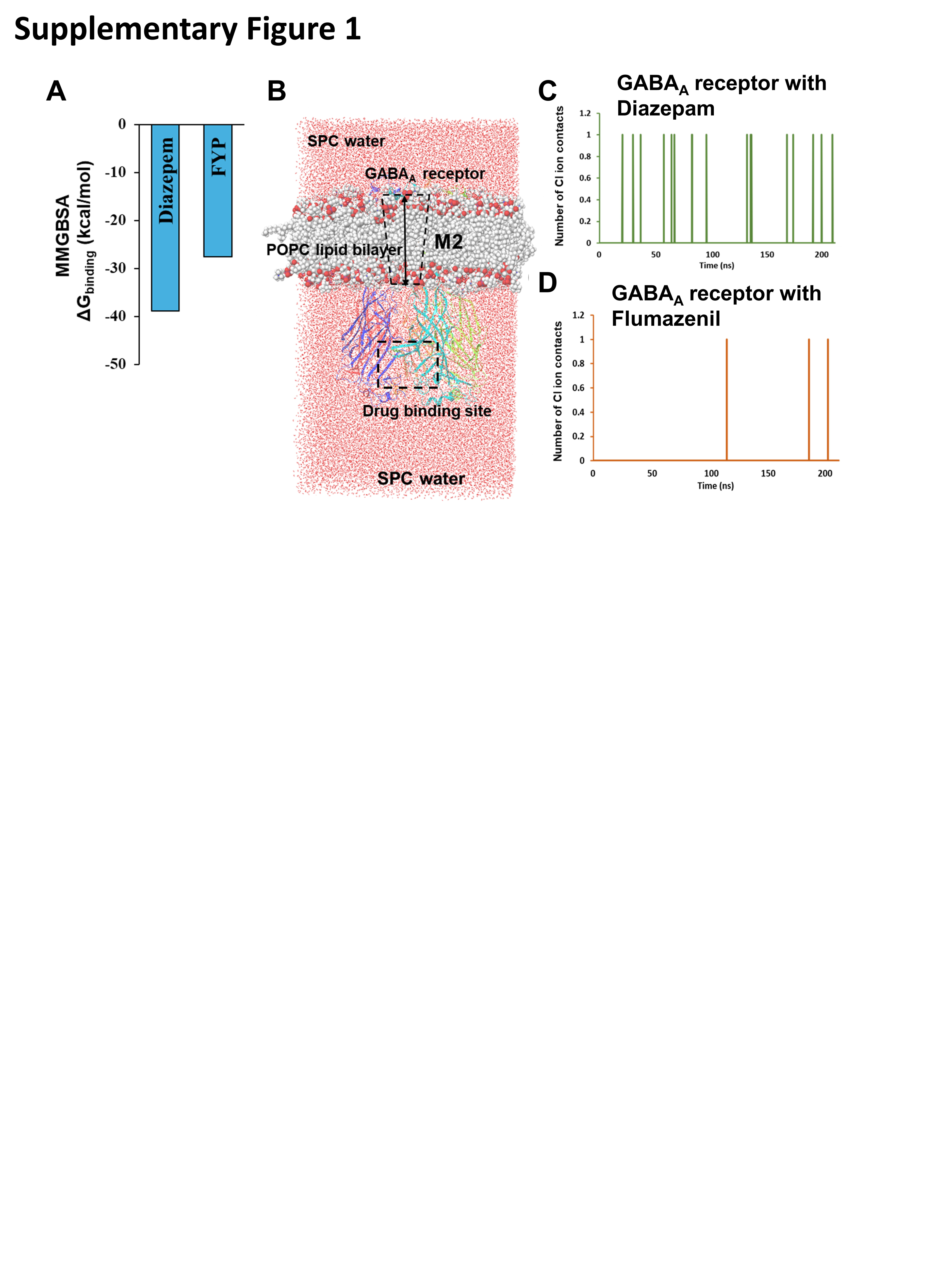

### Supplemental Figure 2

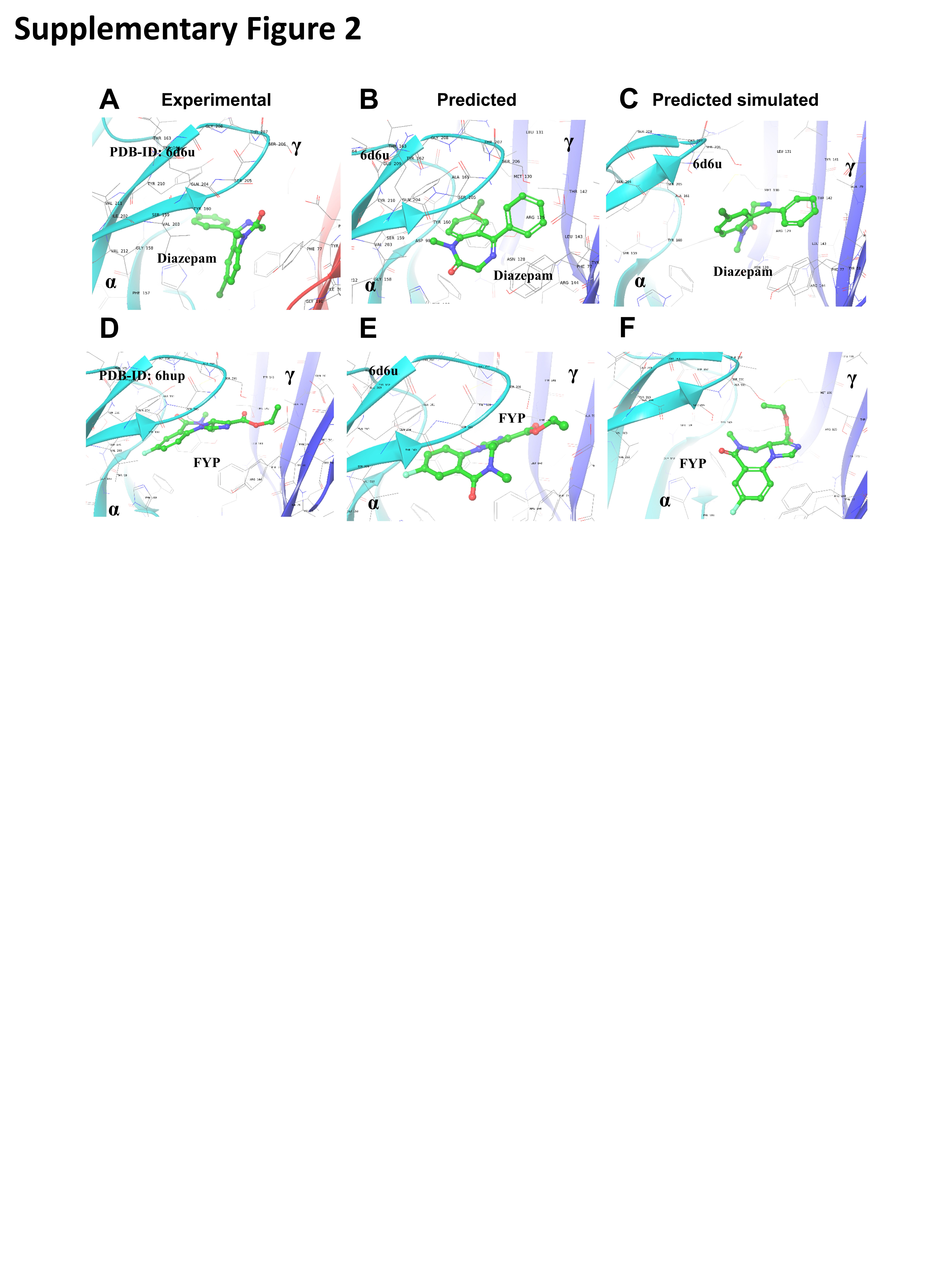

### Supplemental Figure 3

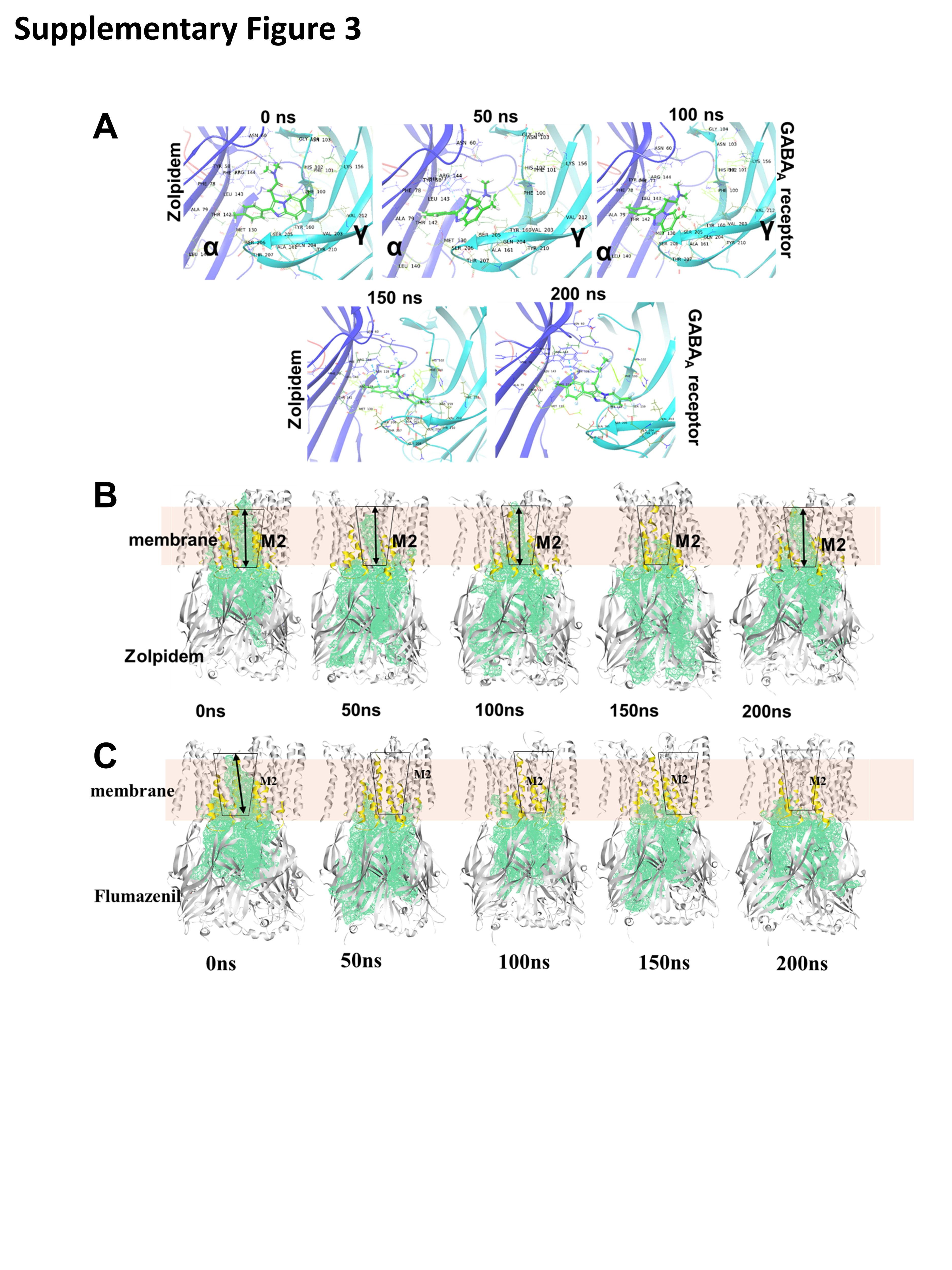

### Supplemental Figure 4

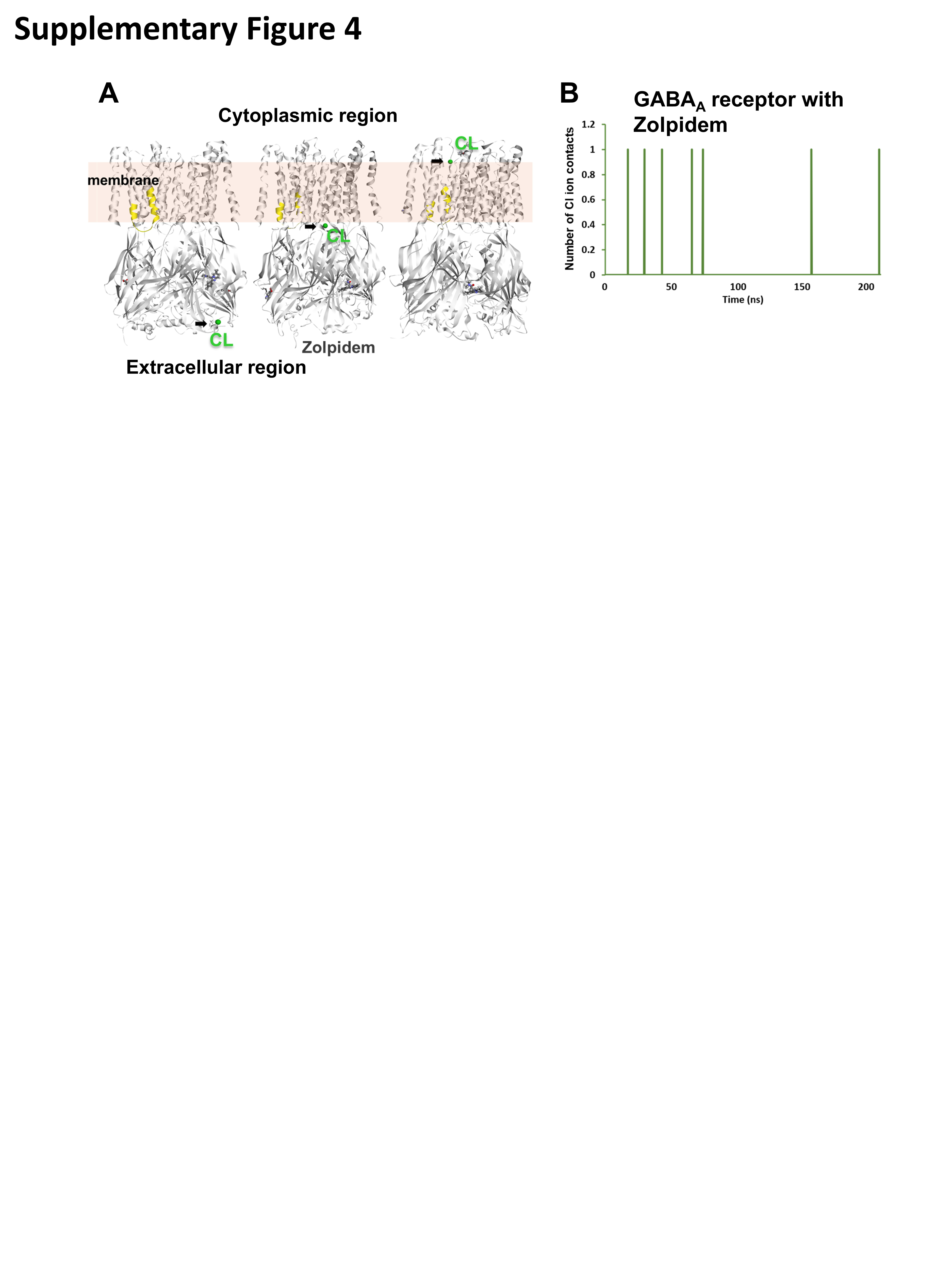

### Supplemental Figure 5

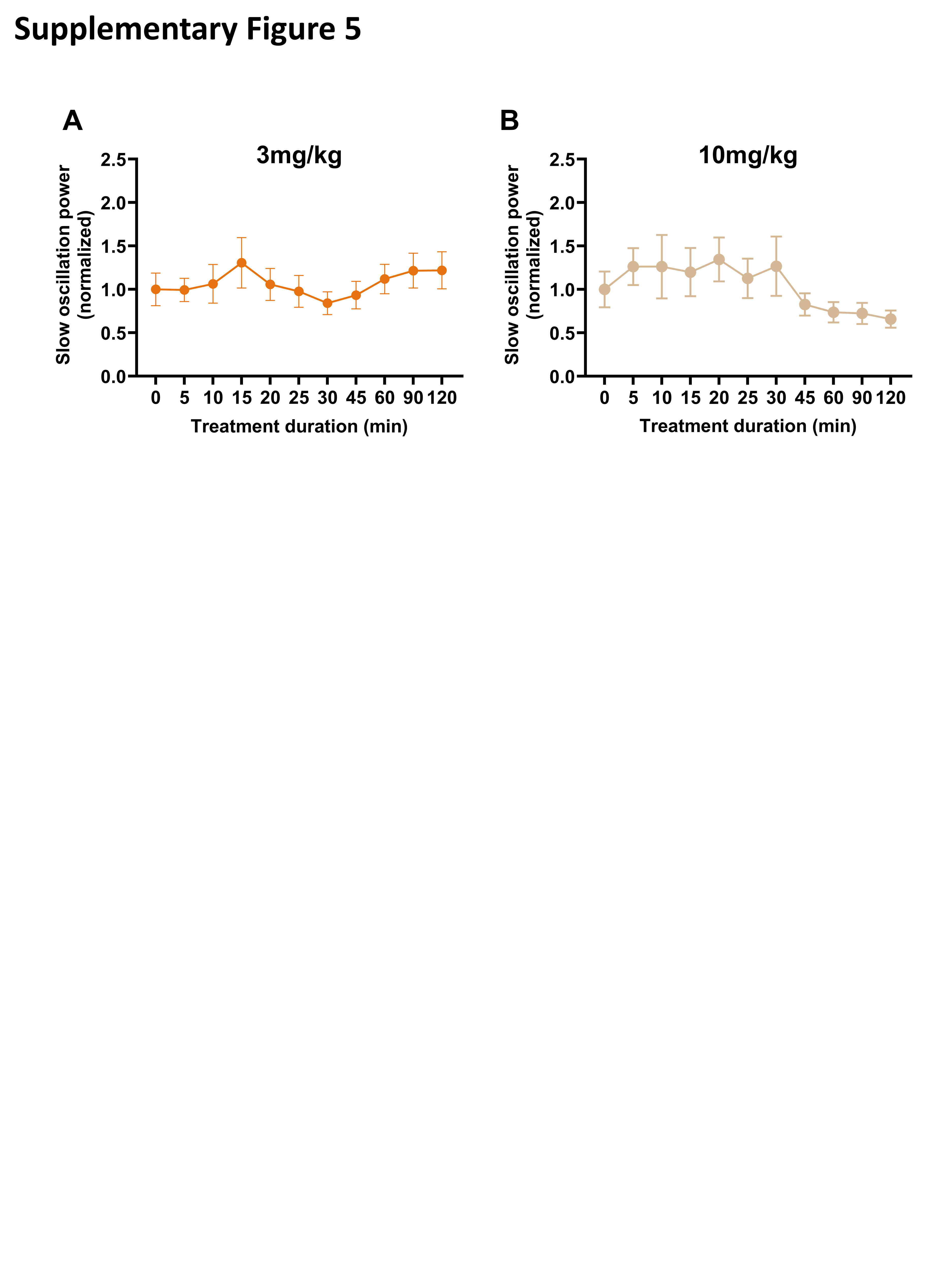

### Supplemental Figure 6

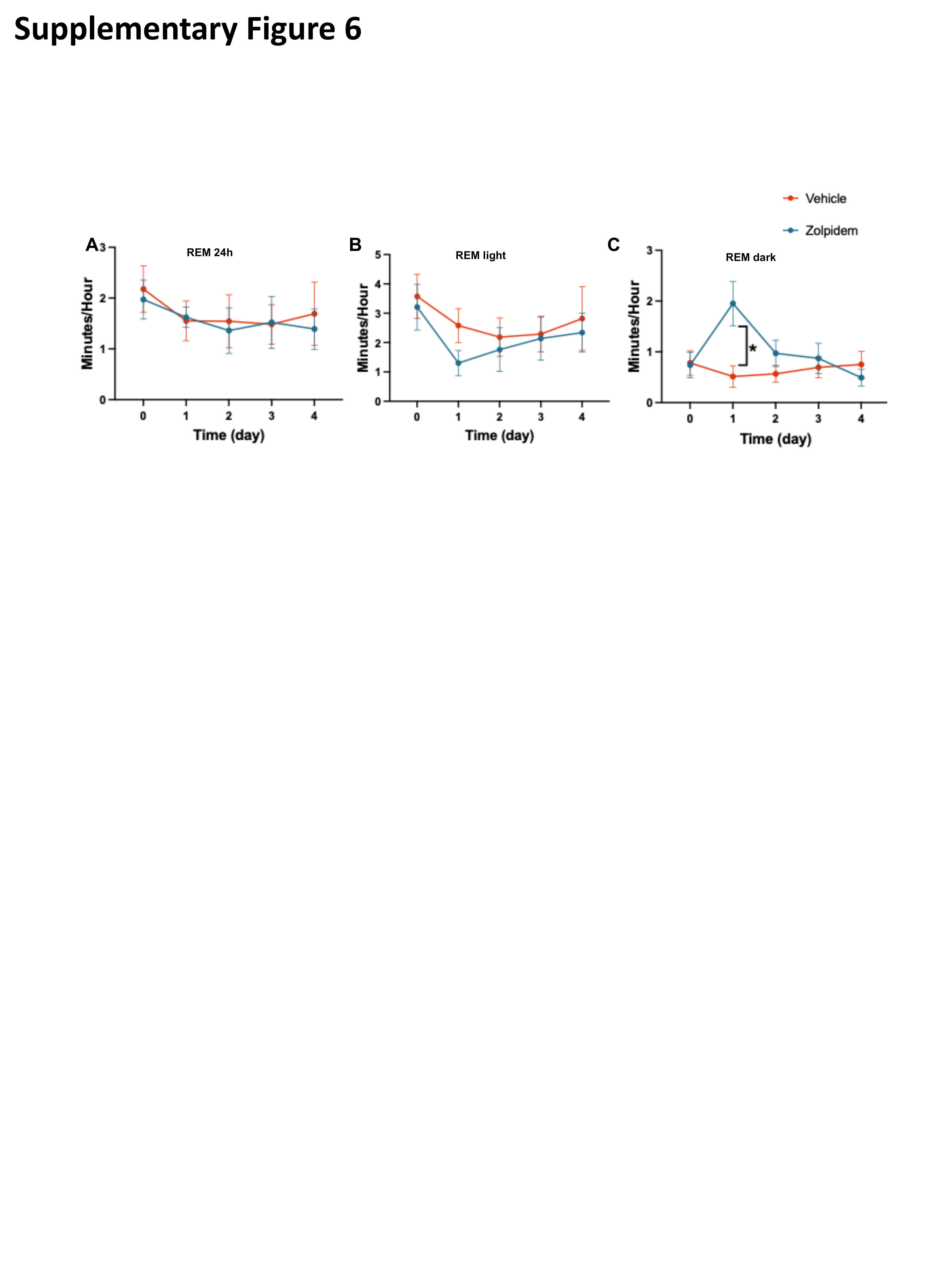
